# A Bactericidal Phospholipase from Archaea

**DOI:** 10.64898/2026.06.10.730825

**Authors:** Chahrazad Taissir, Romain Strock, Yi Wang, Charlotte Guffick, Pauline Misson, Valerie W. C. Soo, Antoine Hocher, Alex Montoya, Pavel V. Shliaha, Hanna M. Oksanen, Jani R. Bolla, Tobias Warnecke

**Affiliations:** Medical Research Council Laboratory of Medical Sciences, London, UK; Institute of Clinical Sciences, Imperial College London, London, UK; Department of Biochemistry, University of Oxford, Oxford, UK; Department of Infectious Disease, Imperial College London, London, UK; Centre for Bacterial Resistance Biology, Imperial College London, London, UK; Department of Genetics, University of Cambridge, Cambridge, UK; Department of Molecular and Integrative Biosciences, Faculty of Biological and Environmental Sciences, University of Helsinki, Helsinki, Finland; Trinity College, Oxford, UK

## Abstract

Archaea kill bacteria, at least on occasion. The molecular underpinnings of these lethal interactions are barely understood. Here, we describe cinquedea, an α/β hydrolase secreted by the halophilic archaeon *Haloferax larsenii* s5a-1. Cinquedea exhibits bactericidal activity in the nanomolar range, killing halophilic *Pontibacillus* bacteria. Bacterial death is accompanied by gross morphological abnormalities, indicative of severe damage to the cell envelope. We predict, and confirm *in vitro*, that cinquedea is a phospholipase, with structural similarities to a phospholipase A1 enzyme isolated from hornet venom. Exposing lipids extracted from a cinquedea-sensitive *Pontibacillus* strain to the enzyme leads to accumulation of lysophosphatidylglycerol, a cleavage product of phospholipase A activity. This is consistent with direct activity of cinquedea against the *Pontibacillus* membrane, which we show is chiefly composed of phosphatidylglycerol. Considered alongside recent findings that some archaea encode bactericidal peptidoglycan hydrolases, these results suggest that archaea can kill bacteria in mechanistically diverse ways. Our work provides a template for future experimental discovery and characterization of bactericidal proteins of archaeal origin and reinforces an emerging view that archaea represent a substantial reservoir for the discovery of new antibacterial compounds.

## INTRODUCTION

Archaea inhabit a wide range of environments where they encounter bacteria, yet the nature of their interactions is poorly understood, especially outside the context of syntrophic communities (Moissl-Eichinger et al. 2018; Strock et al. 2025). Do archaea, at least sometimes, treat bacteria as competitors and actively defend their niche? Might some opportunistically scavenge or even prey on bacteria? If so, what molecular tools do they use to do this?

In bacteria, lethal conflict is common, mediated by a variety of molecular effectors, from large multi-subunit protein complexes like the type VI secretion system to individual proteins (bacteriocins and antimicrobial peptides) to small molecules (Granato et al. 2019). Antagonistic interactions amongst archaea also appear widespread, at least in hypersaline environments (Torreblanca et al. 1994; Atanasova et al. 2013), but the molecular effectors involved in these interactions are known in only a handful of cases (O’Connor and Shand 2002; Kumar et al. 2021; Zachs et al. 2024). One example is halocin H6 from *Haloferax gibbonsii,* a protein that appears to target the Na^+^/H^+^ antiporter of other halophilic archaea, affecting membrane potential and internal pH in a manner that ultimately results in cell lysis (Meseguer et al. 1995).

While conflict is a regular feature of interactions amongst bacteria and amongst archaea, antagonistic interactions *between* members of the two prokaryotic Domains, where archaea kill bacteria or vice versa, are almost entirely undocumented. A small number of studies have explored whether archaea-secreted compounds can inhibit the growth of bacteria, looking for antibacterial activity in cell-free supernatant (CFS), but none of these studies determined a mechanism of action and the bacteria tested were often chosen based on clinical rather than ecological relevance (Shand and Leyva 2008; Atanasova et al. 2013; Ghanmi et al. 2016; Megaw et al. 2019; Castro et al. 2021; Liang et al. 2023).

We recently suggested that antagonistic interactions between archaea and bacteria might, in fact, be common. This proposition was based on the finding that diverse archaea encode homologs of bacterial peptidoglycan hydrolases (Strock et al. 2025). This is curious given that archaea do not encode peptidoglycan, suggesting that these enzymes might be secreted to specifically target bacteria. Indeed, we confirmed experimentally that two peptidoglycan hydrolases from the halophilic archaeon *Halogranum salarium* B-1 exhibit bactericidal activity against the halophilic bacterium *Halalkalibacterium halodurans* (Strock et al. 2025).

Our prior work illustrates that one can sometimes infer archaeal-bacterial conflict by porting insights from the bacterial world – about the proteins that bacteria use to kill other bacteria – to the archaeal world. However, by its nature, such a homology-based approach is limited to discovering tools that are already known to exist as instruments of conflict in bacteria. To find molecular effectors that remain unknown in bacteria or are unique to archaea requires unbiased experimental exploration.

Here, we use such an unbiased experimental approach to dissect one of the few previously reported instances of an archaeon killing a bacterium. We take as our starting point a large-scale screen of antagonistic interactions amongst halophilic prokaryotes. Atanasova and colleagues tested a collection of 68 archaeal and 22 bacterial strains, isolated from different hypersaline environments around the globe, for cross-sensitivity to each other’s secreted compounds (Atanasova et al. 2013). They found 16 instances of cross-Domain sensitivity where supernatant from archaeal cultures inhibited bacterial growth. At least some of these effects were mediated by proteins, as inferred from the observation that proteinase K treatment led to loss of activity. Below, we focus on one of the archaea that produced antibacterial compounds, a *Haloferax* strain designated s5a-1, isolated from the Margherita di Savoia salt pans near Barletta, Italy (Atanasova et al. 2012). Its secreted compounds inhibited the growth of two out of three environmental *Pontibacillus* strains tested in the Atanasova study: SP9-4 and SL-1, isolated from Sedom Ponds (Israel) and a saltern in Sečovlje (Slovenia), respectively. In contrast, the third *Pontibacillus* strain, PV-1 (isolated from a solar saltern in Trapani, Sicily), was unaffected (Atanasova et al. 2013).

## RESULTS

To facilitate identification of the mechanism(s) behind growth inhibition and potentially investigate the genetic basis for differential sensitivity amongst bacterial strains, we sequenced, assembled, and annotated the genomes of the three Pontibacillus strains as well as Haloferax sp. s5a-1 (see Methods, Data S1-8). While these strains had previously been assigned genus-level taxonomic ranks based on 16S rRNA sequencing (Atanasova et al. 2012) whole-genome sequencing affords greater phylogenetic resolution. Using the GTDB toolkit for taxonomic classification (see Methods), we can designate *Haloferax* sp. s5a-1 as *Haloferax larsenii* s5a-1 (Average Nucleotide Identity, ANI = 96.44% to *Haloferax larsenii* JCM13917), consistent with recent independent sequencing and classification of the same strain (Aguirre-Sourrouille et al. 2025). We can further designate *Pontibacillus* sp. PV-1 as *Pontibacillus yanchengensis* PV-1 (ANI = 96.9% to *Pontibacillus yanchengensis* Y32). *Pontibacillus* spp. SP9-4 and SL-1 are almost identical at the nucleotide level (ANI = 99.9%) but substantially divergent from *Pontibacillus yanchengensis* PV-1 (Figure 1A). The closest named species is *Pontibacillus marinus*, but its average nucleotide identity to *Pontibacillus* spp. SP9-4 and SL-1 is only 90.02%. Consequently, *Pontibacillus* spp. SP9-4 and SL-1 should be considered strains belonging to a new species in the genus *Pontibacillus*. Based on the location where the first of these organisms was isolated, we propose to name the two strains *Pontibacillus sedompondensis* SP9-4 and *Pontibacillus sedompondensis* SL-1 and will use these names from hereon in.

**Figure 1.**
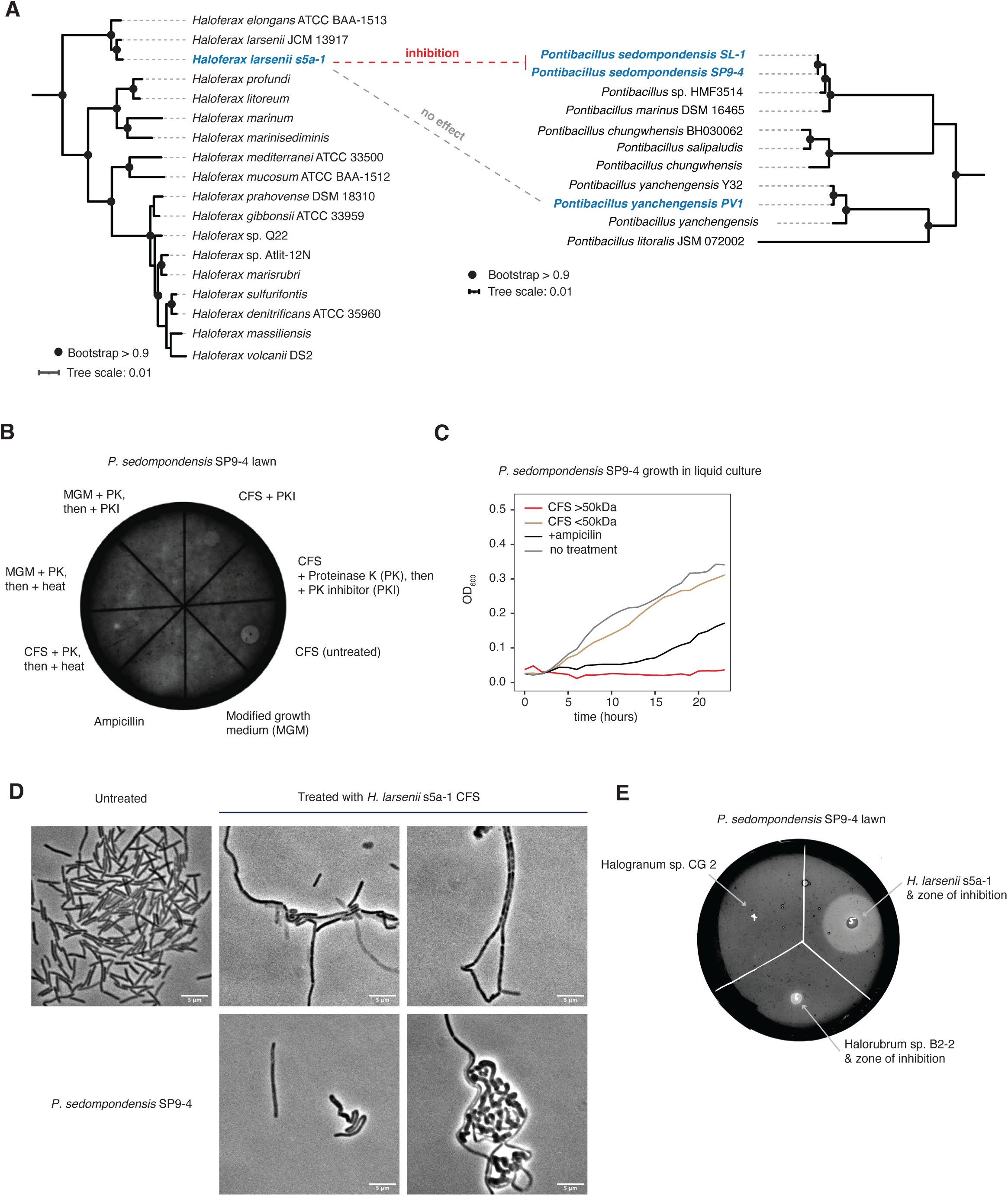
*Haloferax larsenii* s5a-1 secretes a protein with antibacterial activity. (**A**) Phylogenetic placement of the *Haloferax* and *Pontibacillus* strains used in this study. Dashed lines indicate experimentally observed responses to *H. larsenii* s5a-1 supernatant. See Methods for how placement was established. (**B**) Lawn of *P. sedompondensis* SP9-4 treated with *H. larsenii* s5a-1 cell-free supernatant (CFS) along with controls. For samples labelled +PK+PKI, supernatant was first treated with proteinase K (PK) and subsequently with proteinase K inhibitor (PKI), to establish that PK is not itself responsible for antibacterial activity. Minimal growth medium (MGM) and ampicillin (100 µg/ml) served as negative and positive controls, respectively. Note that the zone of inhibition for ampicillin is large and therefore evident as a diffuse, less opaque area. (**C**) Growth of *P. sedompondensis* SP9-4 in liquid culture following treatment with molecular weight-filtered *H. larsenii* s5a-1 CFS. An ampicillin-treated (100 µg/ml) culture was included as a positive control. (**D**) Brightfield microscopy of *P. sedompondensis* SP9-4 cells with and without CFS treatment. Scale bars correspond to 5 µm. (**E**) Live culture spotting assay in which different haloarchaeal cultures were spotted onto a *P. sedompondensis* SP9-4 lawn. Zones of clearing indicate inhibition of SP9-4 growth by *H. larsenii* s5a-1 and, to a lesser extent, *Halorubrum* sp. B2-2 live cultures.

Based on genome annotation and comparison using Prokka and PPanGGOLiN, *P. sedompondensis* SL-1 and SP9-4 are almost identical (with 3,905 and 3,906 protein-coding genes annotated), whereas the *P. yanchengensis* PV-1 genome is significantly larger (4,235 protein-coding genes). This high level of divergence in genome content effectively precludes identification of candidate differences that might underpin differential sensitivity.

### Haloferax larsenii s5a-1 secretes compounds with bactericidal activity

We first sought to reproduce results from earlier work (Atanasova 2013). To that end, we sampled cell-free supernatant (CFS) from *H. larsenii* s5a-1 grown in monoculture and spotted the CFS onto lawns of individual *Pontibacillus* strains (see Methods). Sampling supernatant from across the *H. larsenii* s5a-1 growth cycle revealed that activity against *P. sedompondensis* SP9-4 emerged on day 3 and remained strong into late stationary phase (day 30, Figure S1). For all experiments described below, we use CFS sampled at the onset of stationary phase (day 5).

Zones of inhibition were evident for *P. sedompondensis* SL-1 and SP9-4 (Figure 1B, Figure S2). For unknown reasons, inhibition on plates (but not in liquid culture, see below) was more variable for *P. sedompondensis* SL-1 compared to SP9-4 and we therefore largely focus on *P. sedompondensis* SP9-4 below. We did not observe inhibition for *P. yanchengensis* PV-1, consistent with prior results (Atanasova 2013). In contrast to prior work, we found that treatment of CFS with proteinase K led to loss of inhibitory activity (Figure 1B), suggesting that one or several proteins in the CFS were critical for inhibition. This is further consistent with the observation that, when we passed the supernatant through a 50kDa molecular weight cut-off filter, inhibitory activity was only detected in the high molecular weight fraction retained by the filter (henceforth “top fraction”, Figure 1C). Activity was retained at 60°C but lost at 80°C (Figure S3).

Addition of the CFS top fraction to liquid cultures completely halted growth of *P. sedompondensis* SP9-4 and SL-1 (Figure 1C, Figure S4). *P. sedompondensis* SP9-4 cells treated with the top fraction exhibited pronounced elongation, bulging, and other morphological abnormalities (Figure 1D). Finally, zones of inhibition were also evident after spotting aliquots of growing *H. larsenii* s5a-1onto *P. sedompondensis* SP9-4 lawns, demonstrating direct antagonism in co-culture (Figure 1E).

### Bactericidal activity is caused by an archaeal α/β hydrolase

To identify which protein(s) caused bactericidal activity, we further fractionated the CFS top fraction using size exclusion chromatography (see Methods). The resulting fractionation series was spotted onto plates with *P. sedompondensis* SP9-4 monocultures. Six adjacent fractions that caused inhibition (Figure S5) were processed and analysed using quantitative mass spectrometry (see Methods), along with nine neighbouring fractions that showed no inhibition and therefore contained no or sub-inhibitory concentrations of the protein(s) we are looking to identify. Individual fractions contained peptides from up to 19 proteins, including several proteins not expected to be secreted (e.g. elongation factor EF-2) or to be involved in bactericidal activity (e.g. flagellin), highlighting that CFSs constitute complex mixtures of both actively secreted proteins and abundant cytosolic and membrane-associated proteins (and their degradation products) that are released when cells die (Table S1, Figure S5). To narrow down the list of candidates, we reasoned that the intensity of inhibition should track the abundance of the protein(s) responsible. We therefore compared the phenotypic inhibition profile of the fractionation series with the abundance profile of individual proteins identified across the series. This approach highlighted two high molecular weight proteins whose abundance profile broadly matched the inhibition profile across three biological replicates (Figure 2A): an alpha-amylase family glycosyl hydrolase (pgaptmp_001831_1 in the annotated genome file, Data S1) and an α/β hydrolase containing domain of unknown function DUF726 (pgaptmp_002090_1).

**Figure 2.**
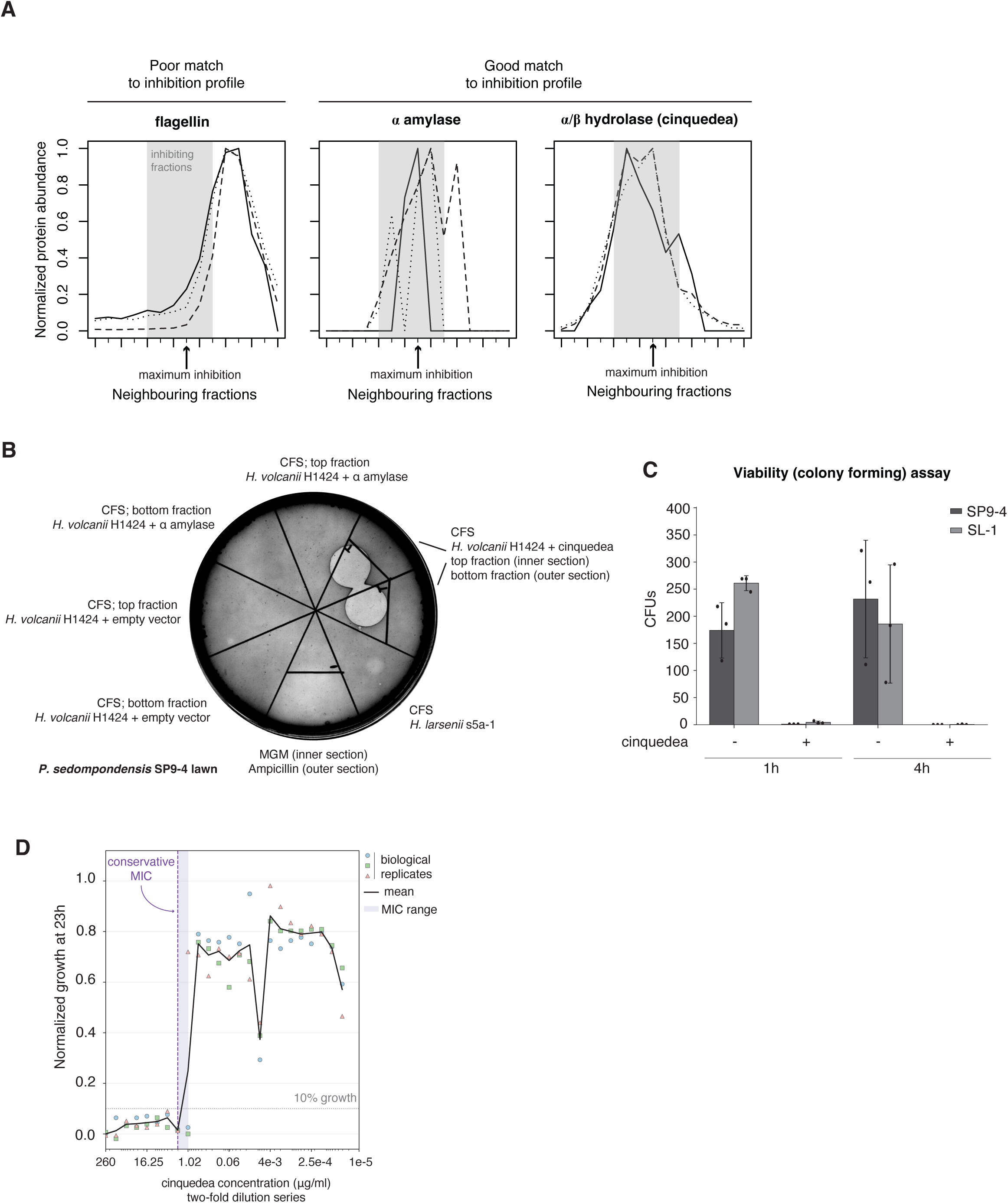
Identification of cinquedea, a bactericidal α/β hydrolase. (**A**) Normalized abundance profiles of three proteins across inhibiting (and neighbouring non-inhibiting) fractions obtained from size exclusion chromatography (see main text). Flagellin is shown as an example of a protein whose abundance profile did not match the inhibition profile well. In contrast, a putative α-amylase and α/β hydrolase (cinquedea) exhibit abundance profiles that match the phenotypic inhibition profile. (**B**) Lawn of *P. sedompondensis* SP9-4 treated with supernatant from *H. volcanii* H1424 heterologously expressing the α/β hydrolase cinquedea, the α-amylase, and various controls. MGM: minimal growth medium. (**C**) Viability assay showing recovery of *P. sedompondensis* SP9-4 and SL-1 cells following treatment with concentrated cinquedea-enriched fractions (see Methods). Liquid cultures (three biological replicates) were exposed to cinquedea or not treated and then left for either 1 h or 4 h before plating. CFU: colony forming units. (**D**) Minimum inhibitory concentration for cinquedea-enriched fractions against *P. sedompondensis* SP9-4. Growth was measured after 23 h across a two-fold dilution series and normalised to the untreated control. The black line tracks mean values across three biological replicates; the shaded region marks the MIC range, with a conservative MIC estimate at 2.03 µg/mL.

Genes encoding these two proteins, both of which are predicted to contain a signal peptide for secretion via the twin-arginine translocation (Tat) pathway, were independently cloned into a plasmid for expression in *Haloferax volcanii* H1424 (see Methods), a chassis we have previously used to test candidate bactericidal proteins from halophilic archaea (Strock et al. 2025). *H. volcanii* H1424 does not encode an ortholog of the α/β hydrolase and CFS from *H. volcanii* H1424 strains carrying either no plasmid or an empty plasmid did not inhibit growth of *P. sedompondensis* SP9-4 (Figure 2B). Neither did the CFS of the strain expressing the putative glycosyl hydrolase. In contrast, strong inhibition zones were evident when *P. sedompondensis* SP9-4 was treated with CFS from strains expressing the α/β hydrolase (Figure 2B). Colony forming assays indicated that the effect of the α/β hydrolase is bactericidal rather than bacteriostatic and morphological observations were consistent with observations from CFS exposure (Figure 2C, Figure S6).

Because of its lethal effect and the fact that its native producer strain *H. larsenii* s5a-1 was isolated from the Margherita di Savoia salt pans in Italy (Atanasova et al. 2012), we name the protein cinquedea, in reference to a short sword used in Italy during the Renaissance period.

We also used CFS from the cinquedea-expressing *H. volcanii* strain to challenge four other Gram-positive halophilic bacteria (*Halalkalibacterium halodurans, Virgibacillus salexigens, Phycicoccus endophyticus,* and *Marinococcus halophilus*) and four halophilic archaea isolated by Atanasova and colleagues (Atanasova et al. 2012): *Halorubrum* sp. E303-2, *Halorubrum* sp. E200-4 *Haloferax* sp. SP10-1 and *H. larsenii* s5a-1 itself. We observed no growth inhibition for any of these species, suggesting that cinquedea-mediated killing is reasonably specific.

To establish the potency of cinquedea, and establish its minimum inhibitory concentration, we carried out broth microdilution assays using preparations of partially purified and concentrated cinquedea against *P. sedompondensis* SP9-4 (see Methods). Using a conservative OD-based inhibition endpoint (∼≤10% fractional growth), the consensus inhibitory concentration across three biological replicates was 2.03 µg/mL (Figure 2D), equivalent to 71 nanomolar for the mature 260 amino acid protein (i.e. after removal of the signal peptide). This is comparable to some well-known bacteriocins such nisin A, mersacidin, and lacticin (Simons et al. 2020). The minimum bactericidal concentration (MBC) was 16.25 µg/mL (0.57 µM), established independently by subculturing 100 µL from selected wells onto agar.

### Cinquedea homologs are predominantly found in archaea

To understand its evolutionary history, we searched for homologs of cinquedea across proteins catalogued in GTDB and UniProtKB (see Methods). Cinquedea homologs are primarily found in archaea (N=479 homologs) and belong to one of two classes: Halobacteria or Nitrososphaeria. Within these classes, cinquedea homologs are common in Nitrosopumilaceae, Nitrosophaeraceae, and Haladaptaceae, with a more patchier distribution in other families (e.g. Haloferacaceae, Haloarculaceae, Figure 3B). Proteins with significant homology to cinquedea are rare in bacteria (N=25) and the genomes where we find them are phylogenetically disparate (Figure 3A) suggesting either sporadic horizontal transfer or contamination of genome assemblies. A majority (82%) of archaeal cinquedea homologs encode a recognizable Pfam domain (69% DUF900, 13% DUF726, 1% cutinase), all of which are associated with the broader Pfam clan covering α/β hydrolases (CL0028). In *H. larsenii* s5a-1, cinquedea is a single-gene transcription unit (Figure 3C) and its genomic context holds no obvious clues as to its functional entanglements.

**Figure 3.**
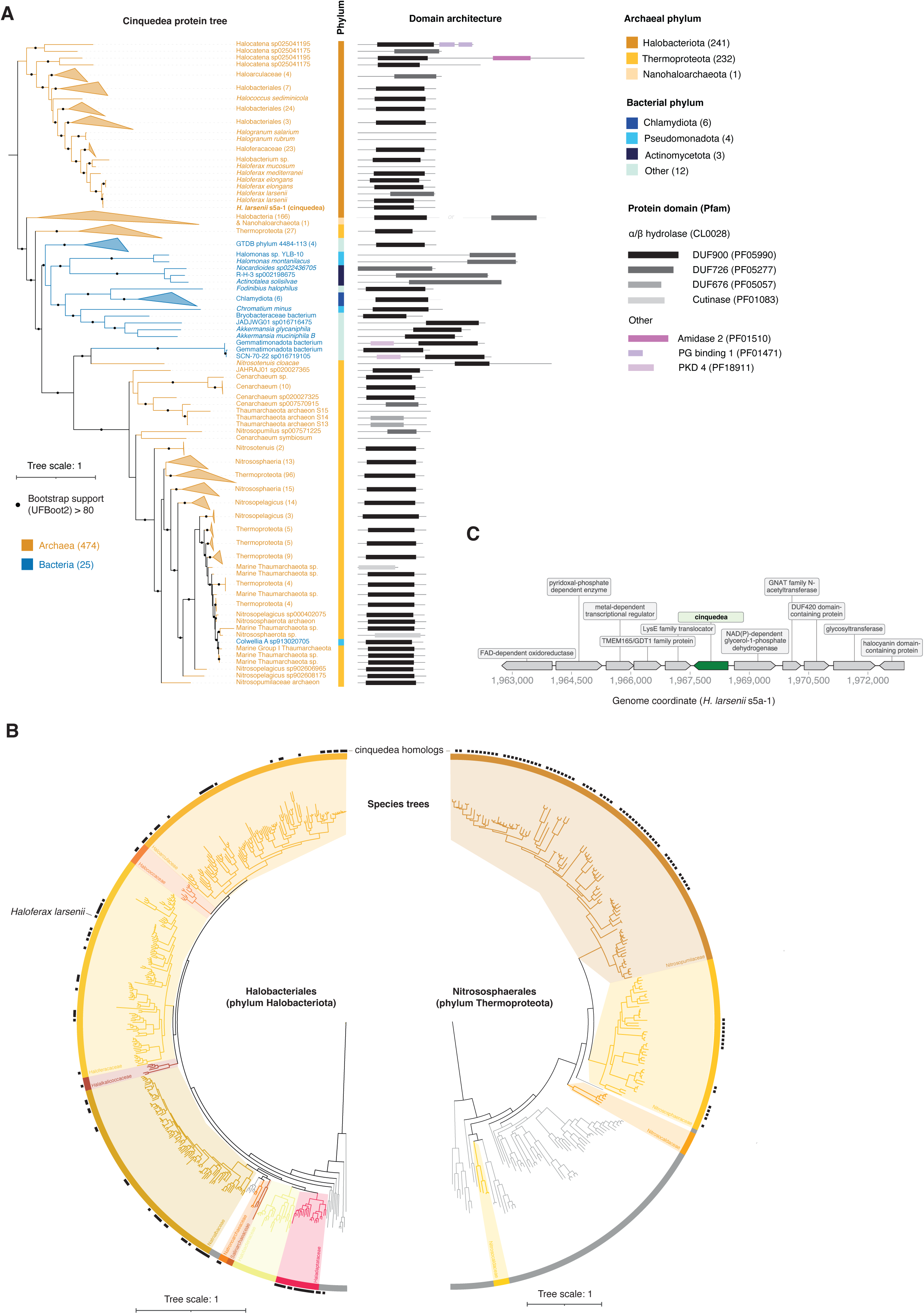
Evolutionary genomic context of cinquedea. (**A**) Maximum-likelihood protein tree of cinquedea homologs identified across archaeal and bacterial genomes. Predicted Pfam domain architectures are shown alongside the tree. See Methods for details of phylogenetic reconstruction. (**B**) Species tree highlighting the distribution of cinquedea homologs across representative Halobacteriales and Nitrososphaerales lineages. Black squares along the outer perimeter indicate genome encoding a cinquedea homolog. Trees are extracted from a larger archaeal phylogeny in GTDB release 214. (**C**) Local neighbourhood (±5kb) of the cinquedea gene in the *H. larsenii* s5a-1 genome.

### Cinquedea has phospholipase activity

α/β hydrolases comprise a set of structurally similar but functionally diverse enzymes, whose substrate preferences are difficult to predict by structural analysis alone (Bauer et al. 2020), including peroxidases, amidases, dehalogenases, and lipases. To narrow down the likely enzymatic activities of cinquedea, we predicted enzyme function using CLEAN, a contrastive learning algorithm (Yu et al. 2023), and searched for structural homologs in the Protein Data Bank (PDB) using DALI (Holm 2022). Both approaches independently associate cinquedea to phospholipase A1 enzymes (EC 3.1.1.32). Interestingly, the closest hit in structural terms is a secreted phospholipase A1 protein isolated from the venom of the black-bellied hornet (*Vespa basalis*) (PDB ID: 4QNN, Figure 4B) which disrupts eukaryotic membrane organization through phospholipids hydrolysis (Ho et al. 1993).

**Figure 4.**
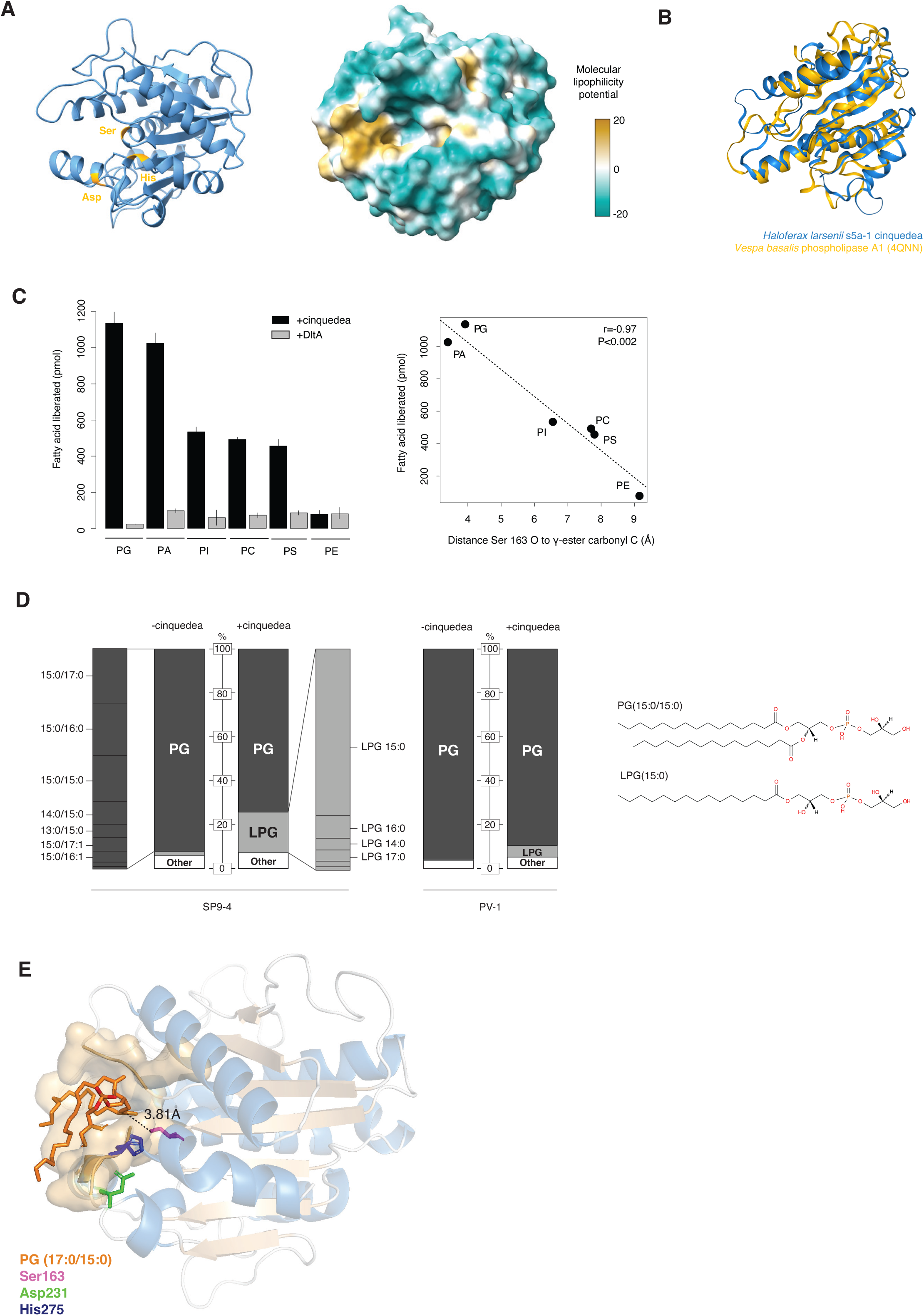
Cinquedea exhibits phospholipase activity. (**A**) Predicted structure of cinquedea highlighting the putative catalytic triad (Ser163, Asp231 and His275) and the molecular lipophilicity potential. The mature protein (without its 30 amino acid N-terminal signal peptide) is shown. (**B**) Foldseek-generated superposition of the predicted structure of cinquedea with that of *Vespa basalis* phospholipase A1 (PDB: 4QNN). (**C**) Left panel: Fatty acid release assay showing hydrolysis of different phospholipid substrates following treatment by cinquedea-enriched fractions. Treatment with purified DltA serves as a negative control. Right panel: correlation between the amount of fatty acid released and the distance between the oxygen of Ser163 oxygen and the γ-ester carbonyl carbon of a given substrate, as determined by in silico molecular docking experiments (see Methods). PG: phosphatidylclycerol (16:0/18:1); PA: phosphatidic acid; PI: phosphatidylinositol; PC: phosphatidylcholine; PS: phosphatidylserine; PE: phosphatidylethanolamine. (**D**) Composition of lipids extracted from *P. sedompondensis* SP9-4 or *P. yanchengensis* PV-1 cultures with and without prior treatment with cinquedea. The most common phosphatidylglycerol (PG) and lysophosphatidylglycerol (LPG) species are indicated. Example structures for PG and LPG are shown on the right. (E) Structural model of PG (17:0/15:0) docked into the cinquedea active site pocket (see Methods for details on the docking procedures). The modelled distance between Ser163 and the ester carbonyl carbon is 3.81 Å.

Cinquedea has a pentapeptide motif (Ser-His-Ser-Leu-Gly) containing the nucleophilic serine, characteristic of known haloarchaeal lipases (X-His-Ser-X-Gly) (Meghwanshi et al. 2022), a well-formed catalytic triad, consistent with lipolytic activity, and a prominent lipophilic cleft suitable for phospholipid binding (see below, Figure 4A).

To establish whether cinquedea does indeed act as a phospholipase, we treated a small collection of phospholipid model substrates with cinquedea purified from our heterologous expression system (see Methods). A protein without known or suspected lipolytic activity (D-alanine--D-alanyl carrier protein ligase, DltA, from *Bacillus subtilis*) was used as a control to establish background rates of fatty acid liberation in the absence of enzymatic activity. We find that phosphatidylglycerol (PG), phosphatidylcholine (PC), phosphatidylserine (PS), phosphatidylinositol (PI) and phosphatidic acid (PA), but not phosphatidylethanolamine (PE) are recognized as substrates by cinquedea, with most fatty acid release observed for PG (Figure 4C). Docking each model lipid into the presumed lipid-binding pocket of cinquedea (see Methods), we find both reasonable enzyme-substrate geometry and, more specifically, that enzymatic activity scales with proximity of the γ-ester carbonyl of the phospholipid to the catalytic serine (Figure 4C).

Next, we asked whether cinquedea can hydrolyse phospholipids present in the membrane of sensitive bacteria. We purified lipids from cinquedea-sensitive (*P. sedompondensis* SP9-4) and -resistant (*P. yanchengensis* PV-1) bacteria, treated the lipid extracts with cinquedea and analysed lipid composition before and after treatment using untargeted LC-MS lipidomics. The lipidomic profile, which we assume chiefly reflects membrane composition, is dominated by different species of phosphatidylglycerol (PG) in both *P. sedompondensis* SP9-4 and *P. yanchengensis* PV-1 (Figure 4D, Table S2). This is not unexpected as PG is also the most prominent membrane lipid in their well-studied cousin *Bacillus subtilis* (Tilburg et al. 2022). PG species in the two *Pontibacillus* strains predominantly have one 15-carbon chain and one carbon chain of variable length (13-18 carbons), reminiscent of the situation in *Staphylococcus aureus*, where the large majority of PG species includes at least one 15-carbon chain, located in the sn-2 position (Subramanian et al. 2023). Lipid extracts from *P. sedompondensis* SP9-4 bacteria exposed to cinquedea exhibit a conspicuous increase in lysophosphatidylglycerol (LPG) 15:0 (Figure 4D), a lipid species that can be derived from PG via cleavage of either the sn-1 or the sn-2 bond by phospholipase A1 and A2 activity, respectively. LPG in general been associated with membrane leakage when present in the membrane at high concentrations (Hull et al. 2004). Accumulation of LPG following cinquedea treatment is much less pronounced for *P. yanchengensis* PV-1 (Figure 4D).

The most abundant PG species in untreated *P. sedompondensis* SP9-4 (PG 28:0 to PG 32:0) were then docked into the cinquedea active site to determine whether they could act as plausible substrates. Focusing on plausible head group orientation, absence of steric clashes, and the distance between the catalytic serine and the γ-ester carbonyl carbon as a readout, we predict that PG is a highly suitable substrate for cinquedea (Figure 4E), with distances in the vicinity or better (3.32-4.08Å) than reported for PA (3.4Å).

These observations further support the notion that cinquedea is a phospholipase A enzyme, directly targeting the bacterial membrane of *P. sedompondensis* SP9-4. Further work will be required to definitively establish whether cinquedea acts as a A1 or A2 type phospholipase and how its activity compromises the *P. sedompondensis* cell envelope. What protects *P. yanchengensis* PV-1 from the action of cinquedea also remains to be determined but it is likely not the absence of target lipids (which are very similar compared to *P. sedompondensis* SP9-4) that determines differential susceptibility. In vivo, differences in the accessibility of cinquedea-targeted lipids, perhaps related to differences in the surrounding peptidoglycan layer, might provide a possible explanation. A similar explanation has been advanced for differential sensitivity of Gram-positive bacteria to human group II A phospholipase A2 (Weiss 2015). This explanation, however, does not explain in vitro observations on lipid extracts, where this barrier should be largely absent.

## DISCUSSION

Archaea encode a variety of lipolytic enzymes, including phospholipases, which have attracted attention for their industrial potential (Feng et al. 2000; Wang et al. 2004; Meghwanshi et al. 2022; Gaonkar et al. 2023; Moopantakath et al. 2023). The physiological and ecological roles of these enzymes, however, typically remain unknown. Our results demonstrate that one of these phospholipases, cinquedea, is bactericidal, killing two closely related Pontibacillus strains.

But did cinquedea evolve to kill bacteria? Several observations, though circumstantial, are at least consistent with this idea: cinquedea is secreted in high abundance (Figure S5) in stationary phase cultures of *H. larsenii* s5a-1, suggesting a) a principal role under conditions where nutrients are (becoming) scarce, and b) that its natural substrate lies outside the cell. There is precedent too for aggressive deployment of lipolytic enzymes, including of type A1 and A2 phospholipases, to compromise membrane integrity, both in the context of interbacterial conflicts, as part of animal venoms, and by the mammalian native immune system to kill pathogenic bacteria (Qu and Lehrer 1998; Laine et al. 2000; Piris-Gimenez et al. 2005; Murakami et al. 2011; Weiss 2015; Flores-Díaz et al. 2016). Finally, the somewhat patchy phylogenetic distribution of cinquedea is not atypical for proteins involved in intermicrobial conflicts. If indeed evolved for bactericidal activity, it is worth noting that this would be a rather nifty evolutionary strategy to target bacteria while avoiding self-targeting, given that archaea employ ether- rather than ester-linked fatty acids (Caforio and Driessen 2017), which should be immune to the action of phospholipase A enzymes.

Yet aggressive deployment against living bacteria is only one of several plausible ecological scenarios. Cinquedea might be secreted to process free/environmental phospholipids, to help in scavenging dead bacteria, for example by breaking open their membranes and making cellular contents available for uptake. It might even act on eukaryotes that inhabit the same niche, like the halophilic green alga *Dunaliella salina,* whose membranes also contains PG, albeit in relatively low abundance (Peeler et al. 1989). Finally, it remains a possibility that cinquedea, through an unknown pathway, is somehow involved in the archaeon’s own lipid metabolism. It is intriguing to note in this context that the gene encoding cinquedea is located next to NAD(P)-dependent glycerol-1-phosphate dehydrogenase, one of the key enzymes in the biogenesis of archaeal membrane phospholipids. The functional significance of this co-localization, however, remains uncertain, especially as it is not broadly conserved across genomes with cinquedea homologs.

To firmly establish the physiological and ecological role of cinquedea it will be imperative to establish its substrate range and specificity, especially focusing on substrates (including living organisms) that *H. larsenii* s5a-1 encounters in its natural environment. Our results imply that, when it comes to bacteria, the realized target range will likely depend not only on the lipids in the bacterial membrane, but also on how accessible these lipids are to cinquedea and therefore on the composition and dynamics of the surrounding peptidoglycan layer. It will also be interesting to determine whether cinquedea-mediated killing of bacteria provides an adaptive benefit to the secreting archaeon, be that by eliminating a competitor for a shared resource or by providing a more direct nutritional benefit. Finally, studying the regulation of cinquedea, for example whether its expression is induced in the presence of (certain) bacteria, would provide further valuable information to understand its physiological role.

Overall, whatever the precise physiological and ecological roles of cinquedea turn out to be, our work, considered alongside previous work (Metcalf et al. 2014; Chen et al. 2024; Strock et al. 2025), highlight that archaea can kill bacteria in mechanistically diverse ways. And the case of cinquedea in particular highlights something else: cinquedea appears to be a predominantly archaeal enzyme, with only a few scattered homologs found outside the archaeal domain. Bactericidal capacity therefore need not be acquired from bacteria via horizontal gene transfer, the likely origin story of archaeal peptidoglycan hydrolases (Strock et al. 2025), but can evolve within the archaeal domain, strengthening the hypothesis that archaea might represent a substantial reservoir for the discovery of new antibacterial compounds (Strock et al. 2025; Torres et al. 2025).

## METHODS

### Archaeal and bacterial strains and growth conditions

All strains were cultivated aerobically at 37°C in a 23% modified growth medium (MGM)(Nuttall and Dyall-Smith 1993). MGM was prepared by diluting artificial seawater (30% w/v) to achieve a final concentration of 23%. The artificial seawater (SW) consisted of 240 g NaCl, 30 g MgCl₂·6H₂O, 35 g MgSO₄·7H₂O, 7 g KCl, 5 mL of 1 M CaCl₂·2H₂O, and 80 mL of 1 M Tris-HCl buffer (pH 7.5) per litre of distilled water. Nutrient supplementation included 5 g peptone (Oxoid) and 1 g Bacto yeast extract (Difco Laboratories) per litre of medium. For solid media and top-agar layers, 14 g/L and 4 g/L of Bacto agar (Difco Laboratories) were added, respectively (Atanasova et al. 2012; Atanasova et al. 2013). To minimize evaporation during incubation, all culture plates were sealed with Parafilm.

### Collection of cell-free supernatant

An inoculum of *H. larsenii* sp. s5a-1 was cultivated in 100 mL of 23% MGM under aerobic conditions at 37 °C with continuous shaking at 180 rpm using an orbital shaker. The incubation proceeded for 120 hours to reach the stationary phase. Post-incubation, cells were removed by centrifugation at 4,500 × g for 10 minutes at 22 °C using an Eppendorf 5810R centrifuge. The supernatant was subsequently filtered through a sterile 0.22 μm syringe filter (28 mm diameter, Sartorius Minisart) to obtain a cell-free supernatant (CFS).

To facilitate downstream protein purification, CFS was concentrated using Vivaspin 20 mL centrifugal concentrator equipped with 50 kDa molecular weight cutoff (MWCO) polyethersulfone membrane (Sartorius).

### Antimicrobial activity assay

Halophilic bacterial strains were cultivated in 5 mL of 23% MGM broth until they reached early exponential phase (OD600 = 0.3-0.5). A 500 μL aliquot of each culture was combined with 3 mL of 18% MGM soft agar maintained at 55 °C. The mixtures were gently vortexed to ensure uniform distribution and then poured onto pre-warmed 20% MGM agar plates. The plates were allowed to solidify for 30 minutes within a laminar flow hood.

Following solidification, 20 μL of the CFS from *H. larsenii* s5a-1 was spotted onto the agar surface of the plate. Negative controls consisted of 23% MGM broth, while ampicillin served as a positive control for antimicrobial activity. The plates were left under the laminar flow hood until the spotted drops had completely dried. Incubation continued until bacterial lawns had developed and zones of inhibition were observable, typically after 1-2 days, at which point plates were imaged.

### Growth assay

To assess the effect of *H. larsenii* s5a-1 CFS on bacterial growth, we used 96-well Nunc black/clear-bottom plates (Thermo Scientific). Bacterial cultures were adjusted to an initial OD600 of 0.05. Each well received 200 μL of a suspension containing a 1:1 by volume mixture of 23% modified growth medium (MGM) and concentrated CFS fractions - either top or bottom fractions obtained using a 50 kDa molecular weight cut-off membrane (MWCO). Plates were incubated at 37 °C in a multi-mode plate reader (FLUOstar Omega) with double-orbital shaking at 700 rpm. Ampicillin was used as a positive control, and 23% MGM without CFS served as a negative control. Bacterial growth was monitored hourly over 24 hours by measuring absorbance at OD600. As an additional control, and to determine whether residual nutrients contributed to observed growth effects, *H. larsenii* s5a-1 cultures were exposed to both fractions of their own CFS. All experiments were performed in triplicate. As done previously (Rogers et al. 2022), to prevent condensation on the plate lids, lids were washed with 10 mL of 10% Triton X-100 in ethanol for 15 seconds, excess solution was discarded, and lids were air-dried in a biosafety cabinet for 30 minutes. Plates were then sealed with Parafilm.

### Live/dead staining and fluorescence microscopy

After 24 h incubation in the 96-well assay, bacterial suspensions were collected and viability was assessed using the LIVE/DEAD BacLight Bacterial Viability Kit (L7012; Invitrogen). SYTO 9 and propidium iodide were mixed 1:1, and 3 µL of the dye mixture was added per 1 mL of cell suspension. Samples were incubated for 15 min in the dark, pelleted by centrifugation (5,000 × g, 5 min), and washed twice with sterile 23% SW. Pellets were resuspended in 23% SW, and 5 µL was mounted on an agarose pad, overlaid with a coverslip, and imaged using a Leica upright fluorescence microscope. Green (SYTO 9) and red (propidium iodide) fluorescence were used to indicate cells with intact versus compromised membranes, respectively.

### Proteinase sensitivity assay

To determine whether the antimicrobial activity of the *H. larsenii* s5a-1 culture supernatant derives from a protein, proteinase K (Invitrogen) was added to the supernatant to a final concentration of 1 mg/mL from a 20 mg/mL stock solution. The mixture was incubated at 50 °C for 1 hour to allow proteolysis. Protease activity was then halted by one of two methods: heating the mixture at 95 °C for 10 minutes, or adding a protease inhibitor cocktail (cOmplete Protease Inhibitor Tablets, Roche) followed by incubation at 37 °C for 1 hour. Treated supernatants were assessed for antimicrobial activity by spotting onto bacterial lawns as described in the antimicrobial activity assay. To ensure assay validity, we included a number of additional controls: (1) CFS treated with protease inhibitor alone to confirm that the inhibitor does not affect the activity of the CFS; (2) MGM supplemented with proteinase K and protease inhibitor to demonstrate that both the enzyme and inhibitor are functional; and (3) MGM with proteinase K followed by heat inactivation to verify that heat effectively halts protease activity.

### Size-exclusion chromatography (SEC)

Cultures of *H. larsenii* s5a-1 were grown to stationary phase and CFS collected and concentrated approximately 100-fold using 20 mL centrifugal concentrators with a 50 kDa MWCO as described above. A 1 mL aliquot of the concentrated CFS was injected into a HiLoad 16/600 Superdex 200 pg column (GE Healthcare) connected to an ÄKTAprime plus chromatography system. Separation was performed at a flow rate of 0.5 mL min⁻¹ and a pressure of 0.88 MPa, using 18% SW as the running buffer. Protein elution was monitored by UV absorbance at 280 nm. Once proteins were detected, fractions were collected at 1 mL intervals. The antibacterial activity of each fraction was evaluated directly after elution, using the top agar antimicrobial assay. Fractions demonstrating activity were pooled for downstream experiments. This procedure was carried out in triplicate with independent biological samples.

### Mass spectrometry - sample processing

Approximately 100 mL of CFS was concentrated to 2 mL using a 50 kDA MWCO filter. 1 mL of this was injected on SEC column and fractionated into 1ml fractions at a flow rate of 0.5 mL/min. For each biological replicate 16 fractions were analysed: 7 fractions before the fraction with most antimicrobial and 8 fractions after. Two hundred µl of every fraction was processed. pH was adjusted to 8.5 with 50mM EPPS, the samples were reduced and alkylated with 10mM TCEP and 20mM cholracetamide for 30 min and then digested with 20ng/µl trypsin (Pierce™ P/N 90059) and 10 ng/µL LysC (WAKO) shaking at 1300 rpm in PCR plates. The digest was desalted using 2mg oasis HLB 30µm resyn packed in a Orochem OF1100 filter plate. Samples were resuspended in 20µl of 0.1% TFA and 5µl of sample was analysed by LC-MS.

### Mass spectrometry - liquid chromatography mass spectrometry (LC-MS) analysis

Chromatographic separation was performed using an Ultimate 3000 RSLC nano liquid chromatography system (Thermo Scientific) coupled to an Orbitrap exploris 240 mass spectrometer (Thermo Scientific) via an EASY-Spray source. Electro-spray nebulisation was achieved by interfacing to Bruker PepSep emitters (PN: PSFSELJ10, 10µm). Peptide solutions were injected directly onto the analytical column (self-packed column, CSH C18 1.7µm beads, 150μm × 35cm) at a working flow rate of 1.25μL/min for 4 minutes. Peptides were then separated using a stepped gradient: 0-45% of buffer B for 66 minutes (composition of buffer A – 95/5%: H2O/DMSO + 0.1% FA, buffer B – 75/20/5% MeCN/H2O/DMSO + 0.1% FA), followed by column conditioning and equilibration. Eluted peptides were analysed by the mass spectrometer in positive polarity using a data-independent acquisition mode as follows: an initial MS1 scan was carried out at 120,000 resolution with an AGC target of 3e6 for a maximum IT of 200ms, m/z range: 409.5-1650. This was followed by sequential MS2 acquisition and fragmentation of ions at 30,000 resolution over 30 variable windows. AGC target set to 3e6 with maximum IT on auto. Normalised collision energy was set to 27%. Total run acquisition time was 80 minutes.

### Mass spectrometry - raw data processing

Data were processed using the Spectronaut software platform (Biognosys, v 18.6.231227.55695) (Bruderer et al. 2015). Pulsar search was performed with default settings for a trypsin/p specific digest with missed cleavage rate set to 3 and a fixed modification of cysteine carbamidomethylation. Variable modifications were allowed for methionine oxidation, protein N-terminal acetylation, asparagine deamidation and cyclisation of glutamine to pyro-glutamate. PSM, Peptide and Protein group identification was carried out an FDR of 0.01. Searches were carried out against the *H. larsenii* s5a-1 predicted proteome and a universal protein contaminants database (Frankenfield et al. 2022) (downloaded 22/01/2024, 381 entries). A mutated decoy database approach was employed with the protein q-value cut-off for the experiment set to 0.01 at the identification level. Quantification was performed at the MS2 level with no value imputation strategy employed.

### Heterologous expression and partial purification

To determine which (if any) of the candidate proteins are responsible for bactericidal activity, we amplified their genes from *H. larsenii* s5a-1 genomic DNA using Q5 High-Fidelity 2X Master Mix (NEB). The primers used (Table S3) contain NdeI and BamHI restriction sites to facilitate cloning into plasmid pTA1392. The amplified genes were verified by electrophoresis on a 1% TAE agarose gel and PCR products excised from the gel and purified using the Monarch DNA Gel Extraction Kit (NEB). Both the purified inserts and pTA1392 plasmid were digested with NdeI and BamHI restriction enzymes (NEBcloner). To prevent vector self-ligation, the plasmid was dephosphorylated using Shrimp Alkaline Phosphatase (NEB). The digested plasmid and inserts were ligated using T4 DNA Ligase (NEB) at a 1:3 vector-to-insert molar ratio for approximately 2 hours, followed by heat inactivation at 65 °C for 10 minutes. The ligation mixtures were then transformed into NEB 5-alpha Competent *E. coli* (High Efficiency) via heat shock transformation. Transformants were selected on LB agar plates supplemented with ampicillin (100 μg/mL). Plasmids from positive clones were extracted using the Monarch Plasmid Miniprep Kit (NEB) and sent for sequencing to confirm successful gene insertion. Confirmed plasmids were then transformed into *H. volcanii* H1424 following the protocol outlined in part 3.3 of (Dattani et al. 2022). As a negative control, pTA1392 without a gene insert was similarly transformed into *H. volcanii* H1424.

Expression of heterologous proteins was induced as follows: transformed *H. volcanii* H1424 cells were consecutively cultured in two different broths, Hv-Ca and Hv-YPC, as described previously (Dattani et al. 2022). A single colony was inoculated into 40 mL of Hv-Ca broth and incubated at 42 °C for two days until the culture reached an OD600 of approximately 0.6. This culture was then used to inoculate 360 mL of Hv-YPC broth supplemented with 1 mM tryptophan (Trp) and incubated at 42 °C with shaking at 180 rpm for 6 hours, reaching an OD600 of around 0.5. Protein expression was induced by adding 36 mL of pre-warmed 25 mM Trp dissolved in 18% SW, and incubation continued under the same conditions for an additional hour (Allers et al. 2010). The CFS was collected by centrifugation and passed through a 0.22 μm syringe filter (28 mm diameter, Sartorius Minisart), then concentrated in a centrifugal concentrator as described above.

Cinquedea was partially purified from CFS by SEC as described before. 1 ml aliquot of concentrated CFS was injected onto a HiLoad 16/600 Superdex 200 pg column connected to an ÄKTAprime plus chromatography system. Separation was performed using 18% artificial seawater as the running buffer at a flow rate of 0.5 mL min⁻¹ and a pressure of 0.88 MPa. Protein elution was monitored by UV absorbance at 280 nm. Active fractions (cinquedea enriched) were pooled and used downstream for Live/Dead Microscopy, MIC and lipidomics.

### Colony-forming unit (CFU) assay

*Pontibacillus* cultures were grown to an OD600 of approximately 0.4. To assess the antimicrobial effect of the CFS from *H. volcanii* H1424 expressing cinquedea, the supernatant was concentrated as before using a Vivaspin centrifugal concentrator with a 50 kDa MWCO PES membrane (Sartorius).

For each bacterial culture, 9 μL of the culture was mixed with 1 μL of the concentrated CFS containing cinquedea. As a negative control, 9 μL of the culture was mixed with 1 μL of the filtrate obtained from the concentrator. Technical triplicates were prepared for both the treatment and control groups. The mixtures were incubated at 37 °C for 1 hour with shaking at 180 rpm.

After incubation, the mixtures were subjected to two consecutive 1:100 serial dilutions. From the final dilution, 100 μL was plated onto pre-warmed 20% MGM agar plates. The plates were incubated at 37 °C for 72 hours. Post-incubation, colonies were imaged using a GelDoc Go Imaging System (Bio-Rad) and quantified using ImageJ.

### Determination of minimum inhibitory concentration (MIC) and minimum bactericidal concentration (MBC)

Determination of MIC and MBC was carried out as per protocol 2a of (Kadeřábková et al. 2024) with minor modifications. MICs of cinquedea preparations were determined by broth microdilution in 96-well plates using 23% MGM as the assay medium. Purified cinquedea stock concentration was quantified prior to the assay as 260 µg/mL total protein.

*P. sedompondensis* SP9-4 cultures were prepared as biological replicates (n = 3). For each replicate, an overnight culture was adjusted to OD600 = 0.08 by dilution in 18% SW to generate the assay inoculum. To estimate the starting inoculum, 10 µL from the positive-growth control well (cells + medium, no cinquedea) was diluted into 10 mL 18% SW (1:1000) and 100 µL plated for colony enumeration.

For the microdilution setup, wells in columns 2–10 were used for the cinquedea dilution series and column 11 was reserved for controls. The plate lid was rinsed with 1× Triton in ethanol (prepared by mixing 1 mL of 10× Triton with 9 mL 99% ethanol), drained, and air-dried prior to incubation. To minimise evaporation, outer wells (columns 1 and 12 and unused perimeter wells) were filled with sterile water, and plates were sealed with parafilm during incubation.

Two-fold serial dilutions of cinquedea were prepared across 24 dilution steps per biological replicate using a continuous vertical dilution scheme (A to H) across three column blocks (columns 2/5/8 for replicate 1, 3/6/9 for replicate 2 and 4/7/10 for replicate 3). Wells were pre-filled with 150 µL 23% MGM, cinquedea stock (300 µL) was added to the first dilution wells (A2/A3/A4; 260 µg/mL) and serial two-fold dilutions were generated by transferring 150 µL sequentially down the columns into 150 µL medium, mixing thoroughly at each step and continuing across column blocks to achieve final concentrations from 260 µg/mL to 3.1×10-5 µg/mL. After the final dilution step, 150 µL were discarded from the last wells to equalise volumes.

Assay wells (cinquedea + medium) and positive growth controls (cells + medium, no cinquedea) were inoculated with 5 µL of the adjusted *P. sedompondensis* SP9-4 inoculum. Negative controls (medium only) received no inoculum. Plates were incubated at 37°C with shaking, and OD600 was recorded hourly for 24 cycles using a microplate reader (Clariostar Plus).

For OD analysis, background was corrected at each timepoint by subtracting the mean OD600 of the medium-only negative controls. MIC was determined for each biological replicate and defined as the lowest cinquedea concentration that yielded no measurable growth relative to the inhibitory control.

To determine the MBC, 100 µL aliquots from or above the MIC wells were plated onto 20% MGM agar and incubated at 37 °C, with a nominal detection limit of ∼10 CFU/mL for undiluted plating. The MBC was defined as the lowest cinquedea concentration at which no colonies were recovered. Inoculum verification plating of the growth-control wells yielded 13–28 colonies following the recommended dilution-and-plating approach, indicating an initial well concentration of approximately 1.3×105 to 2.8×105 CFU/mL.

### Brightfield time-course microscopy

*P. sedompondensis* SP9-4 cultures were grown to exponential or stationary phase and exposed to concentrated recombinant cinquedea. Samples were collected at 0, 2, 4, 6 and 8 h post treatment. Cells were mounted on glass slides with agarose pads and imaged using a Leica DMRB microscope.

### Homology detection

Homologs of the a/b hydrolase cinquedea were identified using MMSeqs2 (Steinegger and Söding 2017) against release 214 of GTDB (Parks et al. 2021) and release 2023_05 of UniProtKB. In both cases, the sensitivity parameter was set to the maximum available in MMSeqs2 (-s 7.5). To remove duplicates, GTDB hits were mapped onto the UniProtKB database using the package “mmseqs map”. Two proteins were considered duplicated if the amino acid coverage between the two proteins exceeded 99%, with perfect identity and no gap along the coverage. In total, this search procedure identified 498 homologs of cinquedea, including 473 archaeal and 25 bacterial proteins.

Protein domains present in a given protein were predicted from Pfam HMM models (release 37) (Mistry et al. 2020) using software HMMER. Gathering thresholds from Pfam were used (--cut_ga option).

### Phylogenetic analysis

Protein trees were constructed as follows. First, protein hits from the homology search were aligned with MAFFT (Katoh and Standley 2013) using the L-INS-I option. Following (Puente-Lelievre et al. 2024) alignment positions with more than 35% gaps were removed using trimAI (Capella-Gutiérrez et al. 2009). Maximum likelihood trees were built using IQ-TREE 2 (Minh et al. 2020) with ModelFinder (Kalyaanamoorthy et al. 2017) (option -m) to automatically select a suitable evolutionary model (selected: “WAG+R8”) and with UFBoot2 (Hoang et al. 2018) (option -B) to generate approximate bootstraps.

The species trees in Figure 3B are subsets of the species tree provided with GTDB release 214 (Parks *et al*., 2022) for orders Halobacteriales and Nitrososphaerales.

Phylogenetic placement for the *Haloferax* and *Pontibacillus* strains was performed with the GTDB toolkit (GTDB-Tk; Chaumeil et al., 2022). First, the classification workflow (command “classify_wf”) was used to find the closest species from the GTDB reference dataset along with ANI values (skani software; Shaw & Yu, 2023). Second, a de-novo tree was constructed (with command “de_novo_wf”) in order to appropriately place the strains SP9-4 and SL-1.

All trees were rendered with iTOL (Letunic and Bork 2024).

### Lipid extraction and treatment

Frozen cell pellets of *P. sedompondensis* SP9-4 and *P. yanchengensis* PV-1 (stored at −80 °C; harvested from 500 mL cultures) were rehydrated in 4 mL of sterile water. Lipids were extracted using a methanol/chloroform method (2:1, v/v) following a modified Bligh and Dyer approach. Briefly, 15mL of cold methanol/chloroform (2:1, v/v) were added to the cell suspension and gently agitated for 15 minutes. The mixture was centrifuged at 3000g for 5 minutes at 4 °C, and the lower chloroform phase (∼3mL) was transferred to a new glass tube. The residual pellet was re-extracted with an additional 2mL of chloroform, vortexed, and centrifuged under the same conditions. The chloroform phases were combined, and 7mL of chloroform and 7mL of 0.2 M KCl were added to facilitate phase separation. The mixture was gently shaken and allowed to separate for ∼3 hours.

The organic phase was divided into 4 glass tubes, corresponding to two treatment conditions for each strain (*P. sedompondensis* SP9-4 and *P. yanchengensis* PV-1): cinquedea treatment and no-enzyme treatment. Nitrogen gas was then used to evaporate the remaining chloroform, and the extracted lipids were retained for enzymatic assays.

For enzyme preparations, 1 mL of purified cinquedea (260µg/mL) was mixed with 1 mL of 18% SW and incubated at 37 °C to form the treatment mixture. The no-enzyme treatment mixture consisted of 2 mL of 18% SW incubated at 37 °C. One millilitre of each mixture was added separately to the lipid extracts from *Pontibacillus yanchengensis* PV-1 and *P. sedompondensis* SP9-4.

Following overnight incubation at 37 °C, reactions were quenched on ice by adding 3mL of cold methanol (3x reaction volume). Phase separation was induced by adding 3mL chloroform followed by 2.7mL of 0.2 M KCl to achieve a final ratio of approximately 1:1:0.9 (CHCl₃:MeOH:H₂O). The lower organic phase was collected and re-extracted then pooled. Samples were dried under a stream of nitrogen gas in amber glass vials, with the headspace flushed with nitrogen before sealing with PTFE-lined caps. The dried samples were stored at −80 °C until further analysis.

### Lipid hydrolysis assay

Phospholipid hydrolysis by recombinant cinquedea was examined using an assay adapted from a previously described method (Imae et al. 2010). Purified recombinant protein was incubated at 37°C for 3 h with lipid preparations containing individual 16:0/18:1 phospholipid substrates: PA, PC, PI, PG, PE, or PS. Following incubation, liberated free fatty acids were quantified using the ACS-ACOD enzymatic method with the NEFA C-Test kit.

### Lipidomics

Extracted lipids were lyophilised and resuspended for reverse-phase liquid chromatography–tandem mass spectrometry (RP-LC–MS/MS) in 68% solvent A and 32% solvent B. Solvent A consisted of acetonitrile:H2O at 60:40 supplemented with 10 mM ammonium formate and 0.1% formic acid, while solvent B consisted of isopropanol at 90:10 with 10 mM ammonium formate and 0.1% formic acid. RP-LC–MS/MS analysis was carried out using a Dionex UltiMate 3000 RSLC Nano system coupled to an Orbitrap Eclipse Tribrid mass spectrometer.

Samples were loaded onto an Acclaim PepMap 100 C18 column, 75 µm × 15 cm, at a flow rate of 300 nL/min. Following a 10 min initial period, solvent B was increased to 65% over 1 min and then to 80% over 6 min. The gradient was maintained at 80% B for 10 min, increased to 99% B over 6 min, and held at 99% B for a further 7 min, giving a total run time of 45 min.

Data were acquired in negative ion mode using data-dependent acquisition. The spray voltage was set to 2.4 kV and the capillary temperature to 320°C. Full-scan MS spectra were collected in the Orbitrap over an m/z range of 300–2000 at a resolution of 120,000. Higher-energy collisional dissociation fragmentation was performed on the most abundant precursor ions within a 3 s cycle time, using a maximum injection time of 50 ms and a stepped normalised collision energy from 25% to 30%. MS/MS spectra were acquired in the Orbitrap at a resolution of 15,000, with a first mass of m/z 75, an isolation window of 1.5 m/z, and a maximum injection time of 50 ms.

The raw data were processed using MS-DIAL 4 (available at https://github.com/systemsomicslab) with MS tolerance of 0.01 Da, MS2 tolerance of 0.015 Da, a minimum peak height of 1000, mass slice width of 0.15 Da and an ms/ms abundance cut off of 200 using a linear weight moving average smoothing with a smoothing level of 3 scans and a minimum peak width of 5 scans. In positive mode, the adduction ion types considered included [M+Na]+, [M+NH4]+, [M+H]+, and [M+H–H2O]+. In negative ion mode, the adduction ion types included [M-H-], [M-H2O-H]-, [M+FA-H]- and [M+Cl]-. Data were searched against in-built MS-DIAL lipid libraries. Only lipids matched at the MS2 level were used in final identification lists. Lipid subclass nomenclature is available at https://systemsomicslab.github.io/compms/msdial/lipidnomenclature.html.

### Molecular docking

Docking was performed using AutoDock Vina (Trott and Olson 2010; Eberhardt et al. 2021). The mature cinquedea model was used as the receptor. Candidate phospholipid ligands were built from SMILES strings, optimised in Avogadro (Hanwell et al. 2012) and converted for docking using Open Babel and Meeko (O’Boyle et al. 2011; Santos-Martins et al. 2025). Vina was used to generate 20 poses per ligand. All poses were manually inspected and we report the best model across all poses, based on the shortest distance between the catalytic serine and Oγ-ester carbonyl carbon, plausible phosphate headgroup contacts, placement of acyl chains in the hydrophobic cleft, and absence of steric clashes.

## CODE & DATA AVAILABILITY

Code and supporting datasets can be found at https://github.com/Chahrazadt87/AvB/.

## ACKNOWLEDGEMENTS

We thank Thorsten Allers for *H. volcanii* strains and plasmids. This work was made possible by core funding from the UKRI Medical Research Council (MC-A658-5TY40) and support from the Department of Biochemistry, University of Oxford (to T.W.), the Royal Society (URF\R1\211567) and a UK Research and Innovation (UKRI) Frontier Research Guarantee Grant (EP/Y036158/1) (to J. R. B.). A.H is supported by a Wellcome Trust CDA (227755/Z/23/Z). H.M.O. was supported by the University of Helsinki and the Research Council of Finland by funding for FINStruct and Instruct Centre FI, part of Biocenter Finland and Instruct-ERIC.

## SUPPLEMENTARY TABLES

**Table S1.** Proteins detected via mass spectrometry in inhibiting and neighbouring fractions. Inhibiting fractions are shown in beige, other fractions in blue. Protein abundance is scaled across fractions so that the fraction where the abundance of a given protein is highest is set to 1. Data for three biological replicates (A-C) are shown.

**Table S2.** Lipidomics of lipid extracts from *P. sedompondensis* SP9-4 and *P. yanchengensis*

PV-1 before and after treatment with cinquedea.

**Table S3.** Primer sequences used in this project.

## SUPPLEMENTARY DATA

**Data S1.** Genome sequence and annotation for the *Haloferax larsenii* s5a-1 largest contig (chromosome) in GenBank format.

**Data S2.** Genome sequence and annotation for the *Haloferax larsenii* s5a-1 second largest contig in GenBank format.

**Data S3.** Genome sequence and annotation for the *Haloferax larsenii* s5a-1 third largest contig in GenBank format.

**Data S4.** Genome sequence and annotation for the *Haloferax larsenii* s5a-1 fourth largest contig in GenBank format.

**Data S5.** Genome sequence and annotation for the *Haloferax larsenii* s5a-1 fifth largest contig in GenBank format.

**Data S6.** Genome sequence and annotation for *Pontibacillus sedompondensis* SP9-4 in GenBank format.

**Data S7.** Genome sequence and annotation for *Pontibacillus sedompondensis* SL-1 in GenBank format.

**Data S8.** Genome sequence and annotation for *Pontibacillus yanchengensis* PV-1 in GenBank format.

---

**Figure S1.**
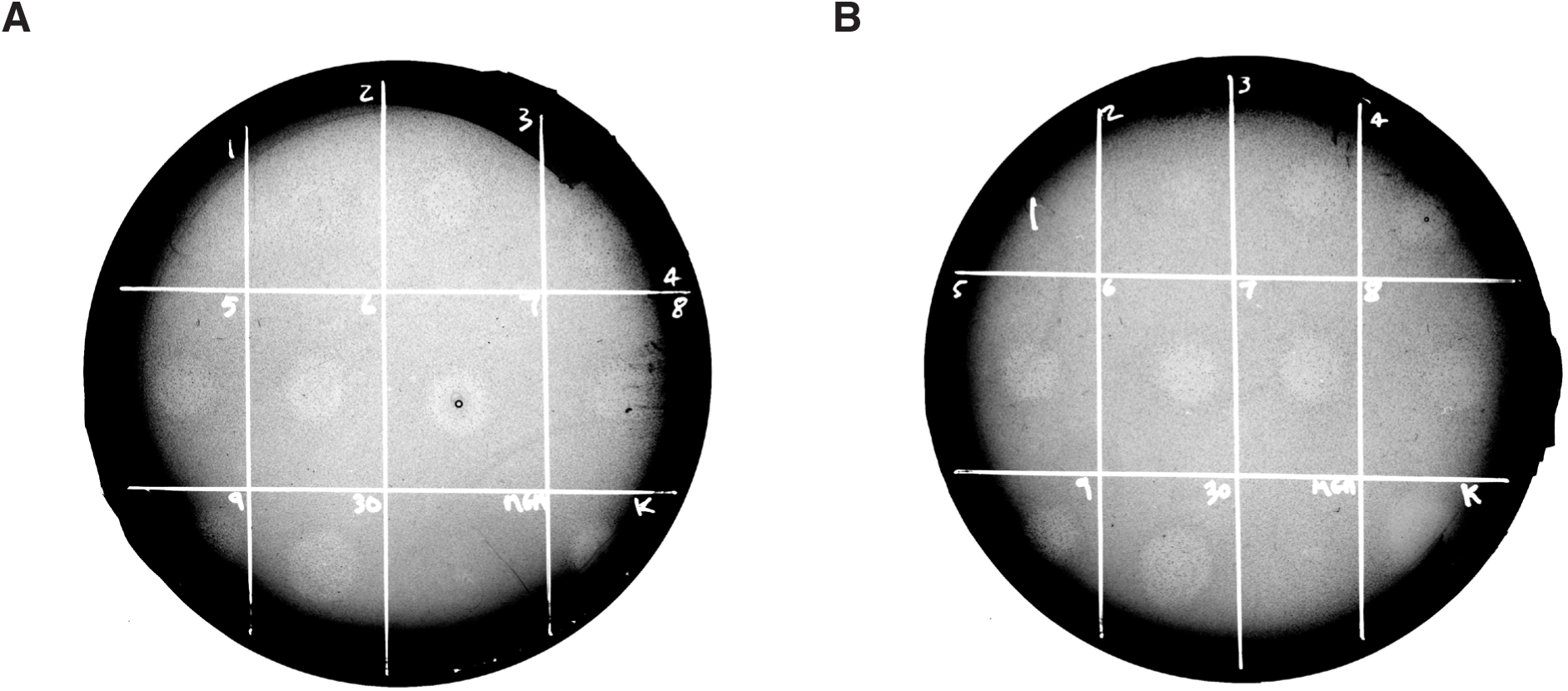
Antibacterial activity of *H. larsenii* s5a-1 CFS across the growth cycle. Spot-on-lawn assays testing CFS collected from *H. larsenii* s5a-1 cultures at different points of growth against *P. sedompondensis* SP9-4 (**A**) and SL-1 (**B**). Numbers correspond to days after *H. larsenii* s5a-1 inoculation at which CFS was collected. Kanamycin (K; 50 µg/ml) and 23% MGM broth serve as positive and negative controls, respectively.

**Figure S2.**
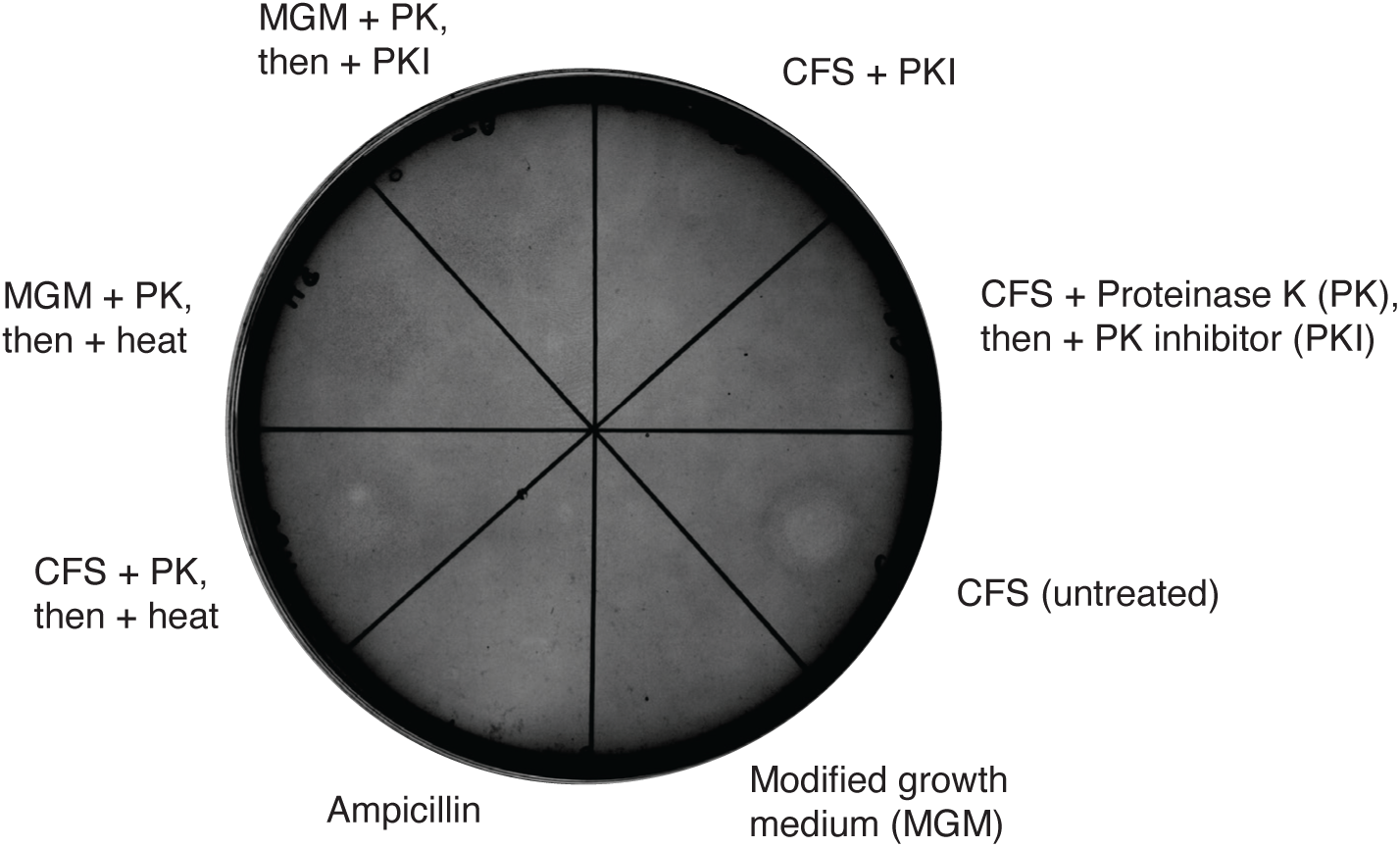
Proteinase K treatment abolishes antibacterial activity of *H. larsenii* s5a-1 CFS against *P. sedompondensis* SL-1. Treatments correspond to those shown in Figure 1B but for *P. sedompondensis* SL-1 instead of SP9-4. For samples labelled +PK+PKI, supernatant was first treated with proteinase K (PK) and subsequently with proteinase K inhibitor (PKI), to establish that PK is not itself responsible for antibacterial activity. Minimal growth medium (MGM) and ampicillin (100 µg/ml) served as a negative and positive controls, respectively.

**Figure S3.**
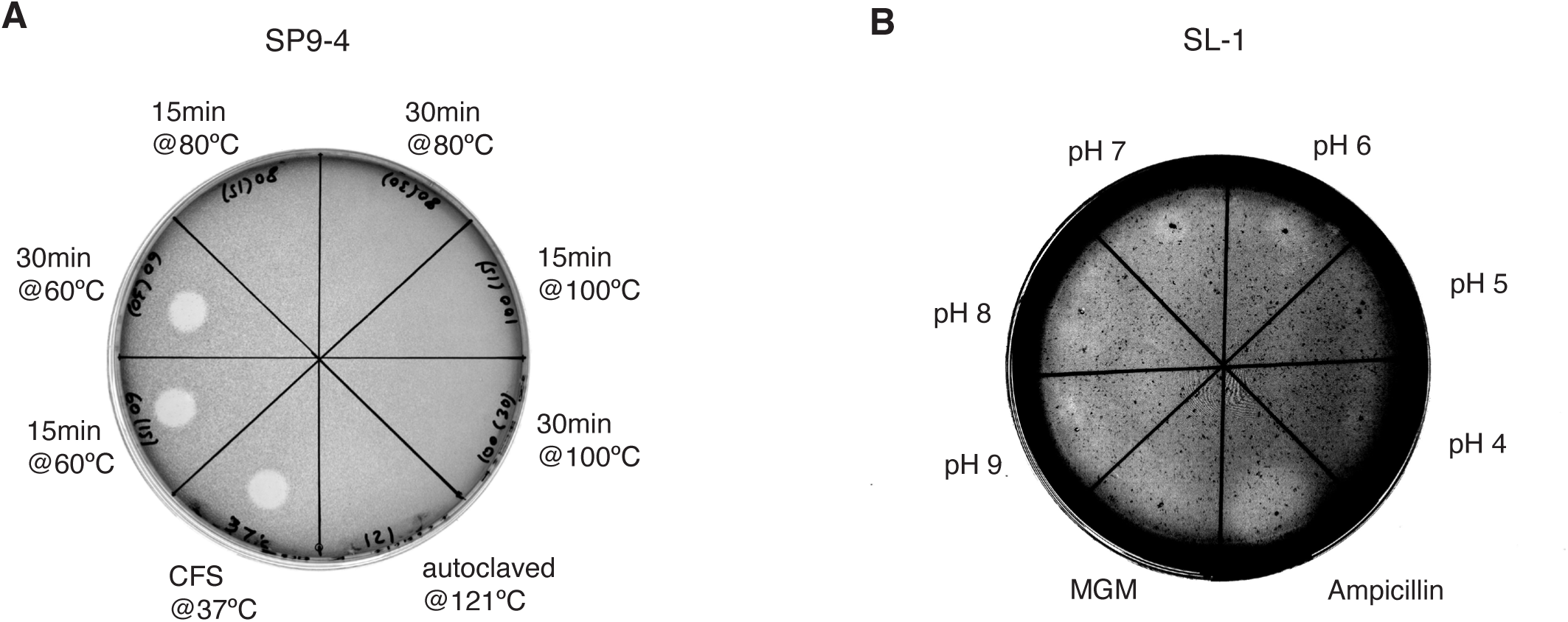
Thermal and pH sensitivity of *H. larsenii* s5a-1 CFS antibacterial activity. (**A**) *H. larsenii* s5a-1 CFS was exposed to the indicated temperature treatments for 15 or 30 min, cooled to room temperature, and tested against a *P. sedompondensis* SP9-4 lawn. (**B**) CFS was adjusted to the indicated pH and tested against a *P. sedompondensis* SL-1 lawn. Clear antibacterial activity was seen at pH 6 and pH 7; inhibition at other pH values was weak or ambiguous. Ampicillin (100 µg/ml) and 23% MGM broth serve as positive and negative controls, respectively. Both conditions used concentrated unfractionated CFS.

**Figure S4.**
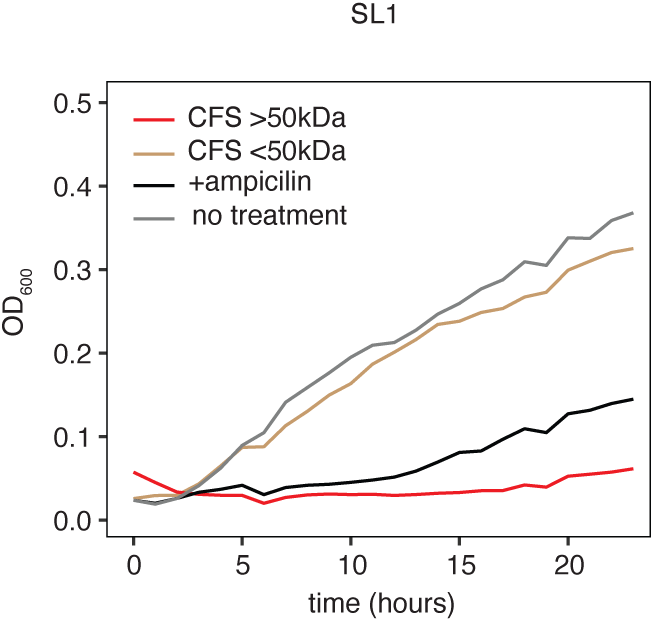
The >50 kDa fraction of *H. larsenii* s5a-1 CFS inhibits *P. sedompondensis* SL-1 growth in liquid culture. The experiments displayed correspond to those shown in Figure 1C but for *P. sedompondensis* SL-1. Growth was monitored for 24 h in 96-well plates after addition of the >50 kDa retentate or <50 kDa filtrate, obtained from *H. larsenii* s5a-1 CFS using a 50 kDa MWCO centrifugal filter. Untreated 23% MGM broth and ampicillin (100 µg/ml) are included controls.

**Figure S5.**
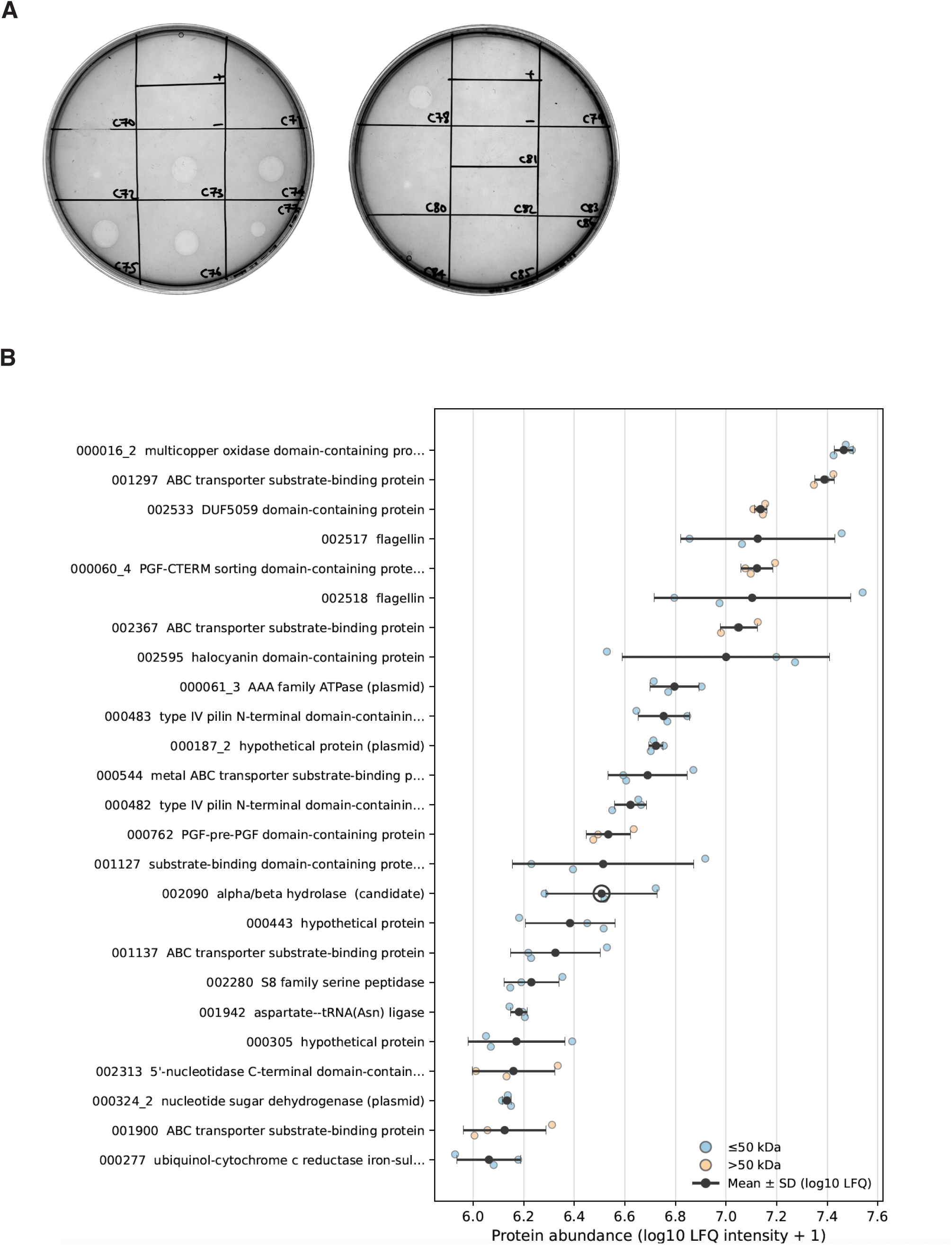
Fractionating the *H. larsenii* s5a-1 secretome. (**A**) Representative spot-on-lawn screening of SEC fractions from *H. larsenii* s5a-1 CFS against *P. sedompondensis* SP9-4. Individual SEC fractions were spotted onto the bacterial lawn, with ampicillin (100 µg/ml) as a positive control and SEC elution buffer (18% sw) as a negative control. Antibacterial activity was restricted to a narrow group of adjacent fractions. (**B**) The 25 most abundant proteins in the day-5 crude *H. larsenii* s5a-1 secretome, before concentration or fractionation, across three biological replicates. Black dots/bars show mean values and their associated standard deviations. Cinquedea, the α/β hydrolase encoded by pgaptmp_002090, is circled.

**Figure S6.**
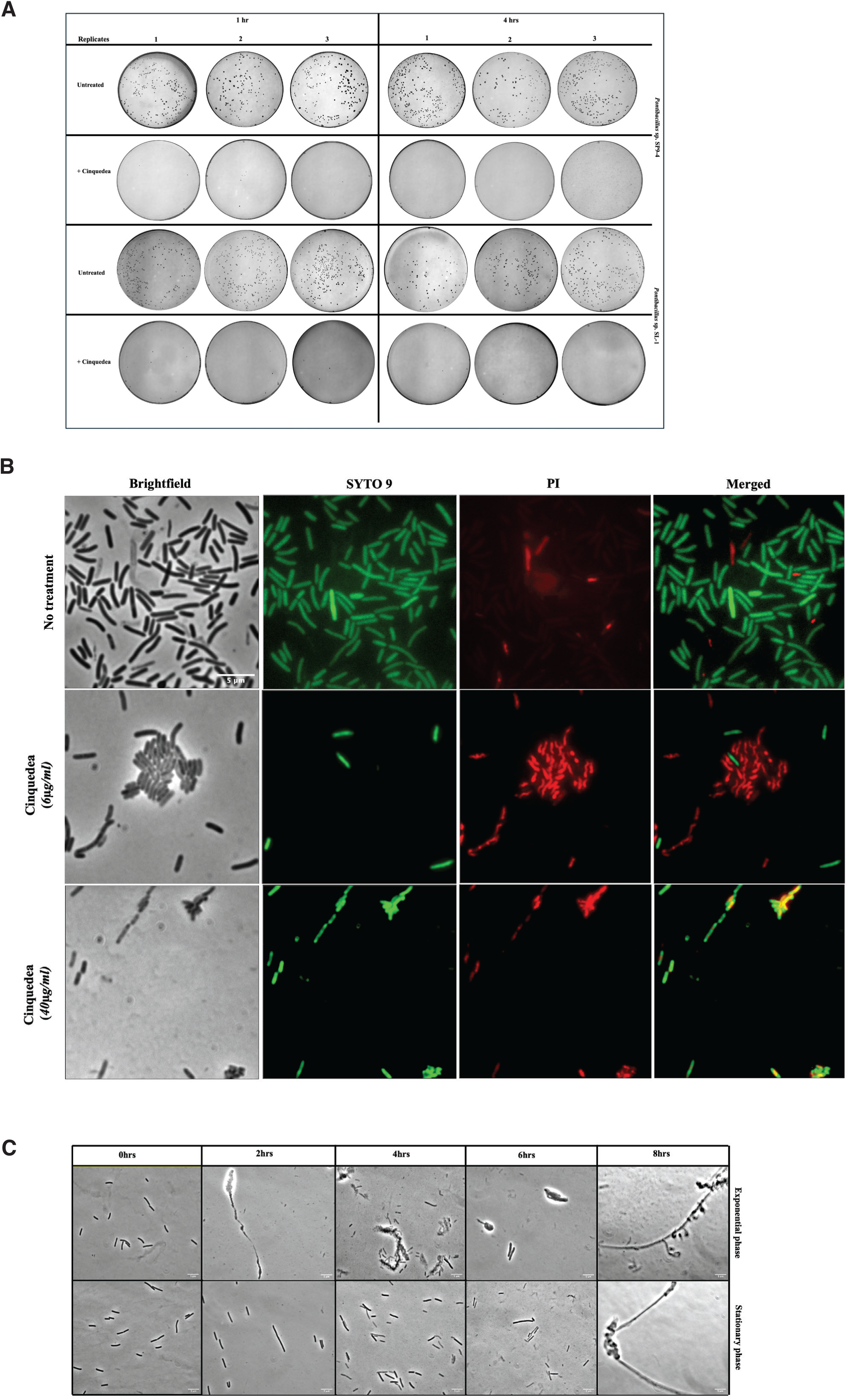
Cinquedea causes loss of viability and time-dependent morphological damage in susceptible *Pontibacillus* strains. (**A**) Dilution plates showing colony recovery from untreated control cultures and cinquedea-treated *P. sedompondensis* SP9-4 and SL-1 cultures after 1 h and 4 h incubation. (**B**) LIVE/DEAD BacLight staining of *P. sedompondensis* SP9-4 after 1 h exposure to purified recombinant cinquedea at 6 µg/ml or 40 µg/ml, compared with untreated cells. Columns show brightfield, SYTO 9, propidium iodide and merged images. Scale bar, 5 µm. (**C**) Time course of *P. sedompondensis* SP9-4 sampled at 0, 2, 4, 6 and 8 h after cinquedea treatment, illustrating progressive elongation or filamentation, localised bulging and the appearance of damaged or lysed cells at later time points.

