## Supplementary material for "A Bactericidal Phospholipase from Archaea": Table S3

**Table S3. Primer sequences used to amplify *cinquedea* (ID 2090) and alpha-amylase family glycosyl hydrolase (ID 1831)**

| Primer pair | Restriction enzyme | Sequence (5’ -> 3’) | Length (bp) | Tm (°C) | GC content (%) |
| --- | --- | --- | --- | --- | --- |
| 2090_F | BamHI | TAGAACTAGTGGATCCGCTGTTGCTTCTCCGTCAGT | 36 | 67 | 50.0 |
| 2090_R | NdeI | GACCTATTGCGCATATGATGTTCCGGTGTGCACAGAA | 37 | 67 | 48.6 |
| 1831_F | NdeI | GACCTATTGCGCATATGCCGACTCGTTACTTCCACGT | 37 | 68 | 51.3 |
| 1831_R | BamHI | TAGAACTAGTGGATCCAACGTCCTCGGTGGTGTTTT | 36 | 66 | 47.2 |
